# MuVEH and mitoMuVEH improve discovery of genetic variation from single cells

**DOI:** 10.1101/2022.11.22.517553

**Authors:** Monica R. Ransom, Krysta L. Engel, Brett M. Stevens, Craig T. Jordan, Austin E. Gillen

## Abstract

Understanding the genetic underpinnings and clonal structure of malignancies at single-cell resolution is critical to accurately predicting drug response and understanding mechanisms of drug resistance and disease evolution in heterogeneous populations of cells. Here, we introduce an accessible, multiplexable, targeted mutation enrichment approach and end-to-end analysis pipeline called MuVEH (Multiplexed Variant Enrichment by Hybridization) that increases the resolution of variant detection in scRNA-seq analysis. When applied specifically to the mitochondrial chromosome (“mitoMuVEH”), this technique can also be used to reconstruct and trace clonal relationships between individual cells. We applied both approaches to two pairs of primary bone marrow specimens from acute myelogenous leukemia (AML) patients collected at diagnosis and after relapse following Venetoclax+Azacitidine (Ven/Aza) therapy. Used together, MuVEH and mitoMuVEH reveal clonal evolution and changing mutational burden in response to treatment at single-cell resolution in these patients. Ultimately, these approaches have the potential to extract additional biological insights from precious patient samples and provide insight into the contributions clonality and genotype have during disease progression.

## Introduction

AML derives from aberrant hematopoietic stem and progenitor cells. As the disease progresses and patients are treated, cells are subject to constant selective pressures leading to the expansion and contraction of clonal populations depending on their fitness. This selection process produces a heterogenous disease, both between patients and within the same patient. This heterogeneity makes AML particularly difficult to treat and is an important driver of relapse in patients [1, 2]. Additionally, this heterogeneity, as well as the phenotypic similarities between healthy and malignant cells, has impeded the characterization of AML biology [3]. Single-cell methods are uniquely positioned to unveil this complexity, which is critical to characterizing this heterogeneity and ultimately developing more effective, targeted therapies.

Current work in the field has begun to approach the questions of somatic mutation profiles as well as clonal lineage tracing from current single-cell datasets. For single-cell somatic variation profiling, many current approaches target the cDNA molecules generated during library construction and amplify specific loci of interest [4–8]. These approaches yield high-resolution results for a small number of genes; however, they require a priori knowledge of the mutational landscape in the patient which is not always available. Similarly, there are two main types of lineage tracing approaches available for single-cell analysis: a modify-and-record approach and mitochondrial lineage tracing. In the modify-and-record methods, a modification (e.g. an expressed traceable tag) is made to an original cell to leave a heritable mark that can be followed through development [9, 10]. While these approaches are powerful and lead to high-resolution clonal maps, they only go back as far as the original labeled cell (i.e. do not capture clonal relationships that exist at the time of labeling) and cannot be applied to intact human cell populations. To trace lineage in humans, work has shifted to tracking naturally occurring background nuclear DNA mutations including SNVs, CNVs, microsatellite repeats, and those found on the mitochondrial chromosome [11–13]. The mitochondrial mutation lineage tracing approach is appealing for several reasons: the mitochondrial genome is large enough to provide a sufficient mutational landscape but small enough to keep sequencing costs low, the mitochondrial genome has increased mutational rates as compared to the nuclear loci [14], mtDNA copy number is high which helps detection in single-cell data, and the mitochondrial RNA or DNA is often captured in the process of making single-cell libraries. There are currently several PCR-based approaches to perform lineage tracing from single-cell libraries, including those utilizing scATACseq [15–17] and scRNA-seq [11] from multiple platforms.

We sought to address some of the issues noted above and increase access to these powerful analyses by expanding our previous hybridization-based single-cell enrichment technique [18] to develop MuVEH and mitoMuVEH (pronounced “movie” and “mito-movie”, respectively). We chose to utilize a hybridization-based approach for four reasons: (1) we can target many (100+) genes with the one pool of probes, (2) we can pool libraries prior to hybridization to save time and reagents, (3) we require less amplification than existing PCR-based strategies, and (4) for ease of execution with this protocol, we can target 64 single-capture libraries in 1 strip tube in about 1.5 days. Additionally, the resulting libraries contain the same indices as the original libraries, which simplifies pooling for resequencing.

These hybridization-based approaches enrich targeted variants and facilitate the accurate measurement of the mutational profile and the clonal relationships of each cell in existing scRNA-seq libraries. Importantly, this protocol functions without modification to the capture or library preparation process. Previous approaches to call mutations from scRNA-seq data have focused on the detection of small numbers of variants in one or a few genes per sample, necessitating upfront knowledge of the driver mutations present in these samples [4, 5, 19]. In contrast, MuVEH uses a panel of probes encompassing more than 100 genes of interest in one hybridization capture (Figure 1A). Similarly, mitoMuVEH uses a panel of probes covering the entire mitochondrial genome to enrich mitochondrial variants that allow us to reconstruct clonal lineages at single-cell resolution. We chose to tile probes across the entire mitochondrial genome based on the observation that single-cell libraries contain both genic and intergenic coverage of the mitochondrial chromosome in our scRNA-seq libraries. We utilize currently available tools to determine mitochondrial variants of interest and use these variants to identify clonal populations of single cells [19, 20]. Both MuVEH mitoMuVEH also take advantage of the indexed nature of the starting material to multiplex 8-10 samples together into one hybridization, cutting down on costs and time. Notably, MuVEH is completely agnostic to the nature of this starting material, final libraries from any single-cell sequencing technique can be used with MuVEH, and the only requirement for mitoMuVEH is capture of mitochondrial RNA or DNA.

**Figure 1.**
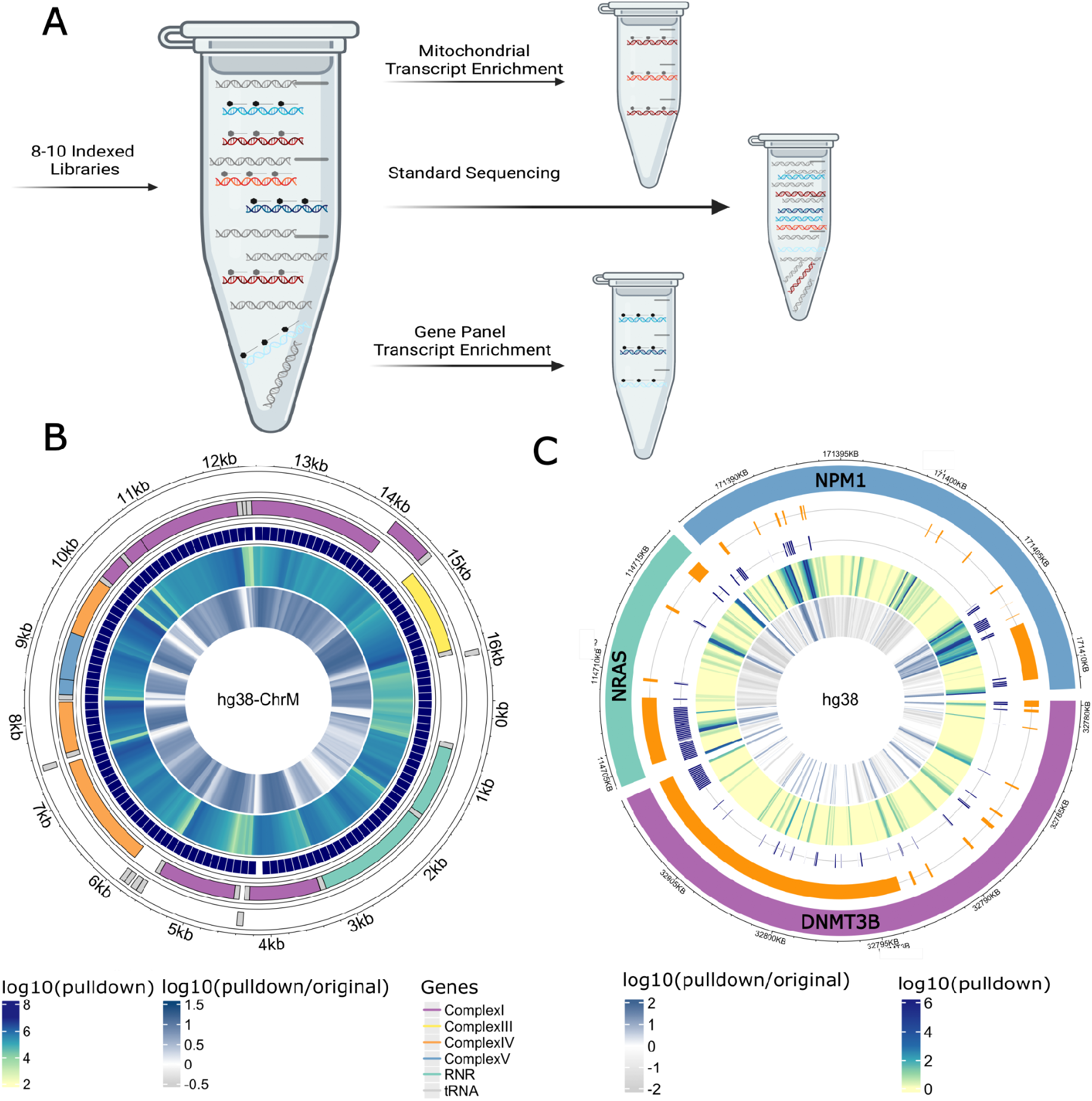
MuVEH and mitoMuVEH enrich for variants in nuclear and mitochondrial RNA. (A) Schematic for hybridization-based enrichment of 10X Genomics final libraries drawn with Biorender. (B) MitoMuVEH enrichment of the mitochondrial genome. Plot from the R package circlize [23] depicting in the rings from outer to inner; mitochondrial location, heavy chain genes, light chain genes, pulldown probe locations, log10(UMI) depth from the pulldown library, and log10(UMI) fold-change between pulldown and original libraries. (C) MuVEH enrichment of 3 genes with high (NPM1), mid (NRAS), and low (DNMT3B) depth in the original 10X libraries. Circle plot depicting in the rings from outer to inner; chromosomal location, gene structure of the full mRNA including all exons, pulldown probe locations, log10(UMIs+1) depth from the pulldown library, and log10(UMIs+1) fold-change between pulldown and original libraries.

Together with our soon-to-be-released end-to-end informatics pipeline, these approaches provide an accessible, multiplexable, and highly flexible set of tools to extract additional data from both existing and future single-cell sequencing experiments. Combined with multimodal single-cell phenotypic measurements including gene expression, surface protein expression, and chromatin accessibility, the genotyping data generated by MuVEH will help investigators better understand the impact of genotype and clonal relationships on the phenotype at single-cell resolution.

## Results

### A hybridization-based method for optimized coverage of nuclear and mitochondrial RNA to yield improved mutation calling from single cell RNAseq libraries

Single-cell RNAseq assays typically have sparse coverage of genes, biases in coverage based on gene location, and technical noise created by PCR amplification [21]. These factors complicate single-cell mitochondrial and nuclear variant identification and render them prone to error. To improve the resolution of this mutation calling we undertook to increase the coverage of both the entirety of mitochondrial variants and targeted nuclear genes of interest to improve variant calling confidence and to help mitigate the dropout effects typically seen in 10X data. We took 10X libraries generated using the 3’ v3.1 protocol and performed enrichment using two custom panels of targets using a tiled probe approach covering the full cDNA of the target gene as well as 5’ and 3’ UTR regions (Figure 1A, methods). Notably, this approach is not limited to 3’ libraries, but also works on 5’ libraries, spatial libraries and could be modified to work with most single-cell approaches. This targeted gene enrichment was originally designed to improve coverage for expression analysis, but we reasoned that it could also be used to improve resolution for mutational analysis.

We designed one panel against the full mitochondrial genome based on the relatively high initial coverage rate in our scRNAseq libraries, which was observed across the mitochondrial DNA independent of gene location. As shown for a representative sample with our mitochondrial pulldown approach, we see high coverage of the full mitochondrial genome with a median depth of 262,562 UMIs per base, and an average fold change of 5.4x in the pulldown compared to the original data (Figure 1B). This is accomplished with approximately 4.5-fold fewer total reads in the pulldown as compared to the original data.

We designed our second panel, an AML-specific gene panel, to target the full-length cDNA and UTR of ∼100 genes that have been shown to be mutated or play a role in AML biology [22]. Figure 1C shows a representative sample of our gene enrichment performance. We see increased coverage of three representative genes with top tertile (NPM1), middle tertile (NRAS), and bottom tertile expression (DNMT3B) in our original dataset. We see an increase in coverage and fold change in the exonic regions targeted across the whole gene body as well as the UTR (Figure 1C). This is accomplished with approximately 5-fold fewer total reads in the pulldown as compared to the original data. Unsurprisingly, this enrichment is limited to regions of the genome that are present in the original libraries and excludes regions far upstream from poly(A) tracts that are not represented in the original libraries. Additionally, some intronic regions see a decreased fold change compared to the original data, due to the lack of probes in this region and these fragments not being resequenced.

### Application of mitoMuVEH and MuVEH to primary AML samples

We applied our hybridization approach to bone marrow mononuclear cells from two AML patients captured at diagnosis and after relapse following Ven/Aza-mediated remission. For these 4 samples, we first called mitochondrial mutations from the single cell data treated as bulk on the diagnosis and relapse samples separately. Bulk SNV calls for each timepoint as well as for a combined diagnosis + relapse were generated. These variant lists were combined and filtered using MQUAD [19], which uses a binomial mixture model to identify variants that are informative for determining clones. These filtered variants were then used as input into cellsnp-lite [20] to call mutations in single-cell mode, ultimately producing matrices of variant counts per cell. We then used vireoSNP [24] to assign probability estimates to the clones and visualized them using Seurat [25]. Transcriptome and CITE-seq analysis of 22 target proteins were combined and used to generate a UMAP projection colored by cell type (Figure 2A). VireoSNP mitochondrial clone assignments were then used to generate alluvial plots (Fig 2B-C) and projected on the transcriptome UMAP, split by timepoint, and colored by mitoMuVEH-defined clone (Figure2D-E). For each diagnosis/relapse pair, six mitochondrial clones were confidently identified based on a plateau in the ELBO score reported by vireoSNP. In Patient 1 at diagnosis, there are two main clones, M1 and M2, which label the monocytic and primitive cells respectively (Figure 2D). Upon relapse, both of these clones collapsed and a phenotypically distinct myeloid-committed population of clones, M3-6 expanded (Figure 2B). In contrast, the primitive cells in Patient 2 at diagnosis are indistinguishable from putatively normal lymphocytes (clone M1), likely due to a lack of differentiating mitochondrial variants in the primitive AML population (Figure 2E). Like Patient 1; however, this patient also experiences a complete loss of phenotypically primitive AML cells at relapse, concurrent with an expansion of subclonal, phenotypically monocytic cells(Figure 2C). These results are consistent with previous studies on AML heterogeneity post-relapse.

**Figure 2.**
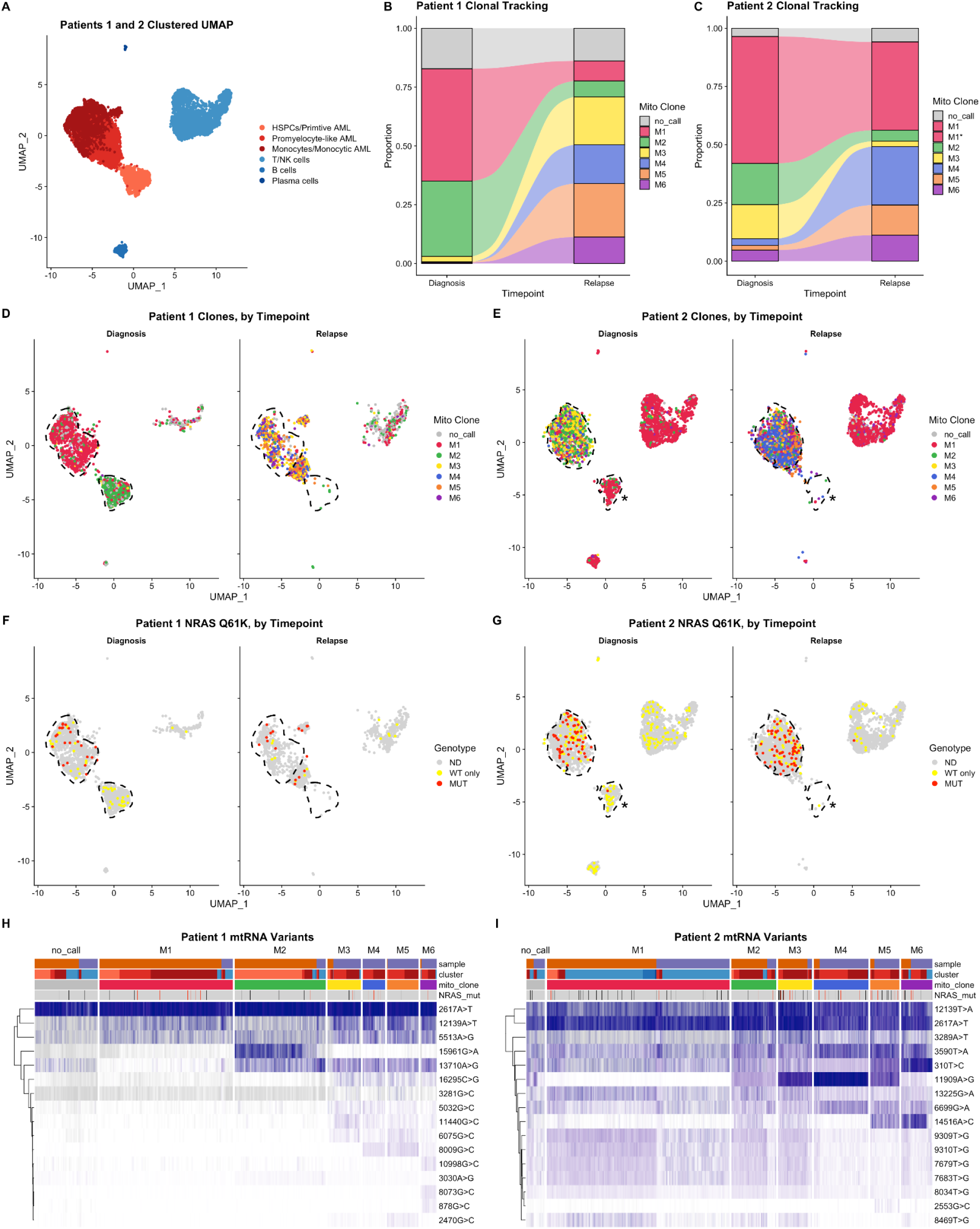
MuVEH and mtMuVEH can be used to infer clonal lineage and mutational status in AML diagnosis and relapse samples. Here, bone marrow mononuclear cells from two AML patients captured at diagnosis and after relapse following Ven/Aza-mediated remission are interrogated using CITE-seq with MuVEH and mitoMuVEH targeted enrichments. (A) A combined clustered UMAP projection based on the transcriptome and surface proteome (22 protein targets). (B, C) Alluvial plots show the expansion and loss of clones following relapse. (D, E) A UMAP projection of the clonal structure of the two patients at diagnosis and relapse. (F, G) UMAP projection colored by the mutational status of a prognostic driver mutation NRAS Q61K variant observed in both patients for diagnosis and relapse samples, the variant allele is shown in red, the reference allele in yellow, and no call is shown as gray. (H, I) Heatmap projections showing the proportion of mitochondrial variants of interest per cell. The heatmap is also annotated by cluster designation based on the transcriptome and surface proteome as well as for the presence of the driver NRAS mutation in these samples.

Both Patient 1 and Patient 2 had a known upfront driver mutation in NRAS at position Q61. A UMAP was generated for each sample using the combined transcriptome and proteome cluster assignments. MuVEH was then used to visualize this diagnostic driver mutation in both patients (Figure 2F-G). In both patients, the variant allele (red) is restricted primarily to the monocytic AML populations, with the detection of only the WT allele (yellow) in the most primitive AML cells and normal (non-neoplastic) lymphocytes.

MuVEH and mitoMuVEH together revealed complex clonal structures in both patients by leveraging mtRNA and nuclear RNA SNVs. These can be visualized together on heatmaps showing the clone informative mitochondrial variants per cell colored by their expression(Figure 2H-I). Further annotation of these cells by the cluster designation, mitochondrial clone call, as well as SNV mutations of interest help uncover the clonal complexity in these samples.

## Discussion

It is often difficult to determine the mutational profile or clonal lineage in patient-derived cells. Here we demonstrate a method to enhance the coverage of genomic regions to aid the identification and quantitation of these variants. Requiring minimal enrichment steps, our protocol is able to enrich these loci in a multiplexed and efficient way. This protocol can target mutations in arbitrary numbers of genes, as long as there is sufficient starting material for the combined genes in the initial library, and across the entire mitochondrial genome. Our analysis shows that this technique can be used to increase detection of mutations in primary human samples, and is particularly well suited to longitudinal study designs as it can track clonal evolution across time.

The strength of this approach is the use of a hybridization technique developed previously by our group [18] and implemented in a 10X Genomics protocol to target the enrichment of genes of interest in previously prepared libraries. We used this approach to target many genes known to be mutated in AML patients and the full mitochondrial genome. These libraries are highly enriched for the targets and can be sequenced at a relatively low depth and still yield high coverage in our regions of interest. The number of genes targeted was relatively low in our case (110 genes) but performance could be improved further with an increased number of genes. This approach allows for a less biased interrogation of the mutational landscape in single-cell analysis when compared to existing methods.

With the possibilities enabled by this approach, we hope to further elucidate the role that genotype plays in cancer formation, progression, and resistance to treatment. The method presented here is not specific to our cell type, disease, or even single-cell sequencing technique, and can be used to help increase the knowledge gained from precious samples. For example, we MuVEH to be particularly useful as a tool to enhance long read single-cell sequencing protocols. As this field is evolving, we envision this approach can be utilized and improved to study a wide variety of important biological questions.

## Methods

### 10X targeted custom AML mutation panel

A 10X targeted custom panel was designed using the 10X Genomics custom panel design software. 111 human gene targets were input into the software yielding a design of 5475 baits, which were then analyzed with 10X Targeted Depth tool for percent on target in our previously sequenced libraries (ranging from 0.13-0.28%). A percent on target of greater than 0.1% is needed for optimal hybridization efficiency, therefore our panel passed the criteria to proceed. These baits were ordered from IDT as biotinylated probes in a 16-reaction Discovery Pool and treated as per the manufacturer’s specs.

### 10X targeted custom Mitochondrial DNA panel

The full mitochondrial genomic DNA sequence was used to design a tiled mitochondrial 10X Genomic custom targeting panel. This panel design yielded 137 probes that were ordered from IDT as biotinylated probes pooled in a 16-reaction xGen Custom Hyb panel. The percent on target of this panel was measured as described above and exceeded the 0.1% needed for hybridization efficiency.

### Standard 10X targeted libraries

Standard 10X targeted libraries were made as per the protocol from 10X Genomics using either the AML mutation panel probeset or the mitochondrial probe set. Briefly, libraries were measured by qubit for concentration, and libraries that had less than 300ng of material were amplified. Eight to ten libraries with compatible indexes were pooled with 300ng of input each. Samples were lyophilized with Cot DNA and Universal blockers. Libraries were then hybridized with the custom panel following the fully custom panel guidelines using 2 μl of the probe pool. Libraries were captured with Dynabeads M-270 Streptavadin (ThermoFisher cat # 65305) beads and washed 4x with the provided wash buffer. Libraries were amplified for 13 cycles of PCR for the AML panel and 11 cycles for the mitochondrial panel. The resulting libraries were normalized to 4nM for sequencing.

### scRNA-seq data processing

Raw sequencing data for gene expression, antibody-derived tag (ADT; surface protein), and hashing libraries were processed using STARsolo 2.7.8a [26] with the 10X Genomics GRCh38/GENCODE v32 genome and transcriptome reference version GRCh38_2020A; https://support.10xgenomics.com/single-cell-gene-expression/software/release-notes/build#GRCh38_2020A or a TotalSeq barcode reference, as appropriate. Hashed samples were demultiplexed using GMM-Demux [27]. Next, cell-containing droplets were identified using dropkick 1.2.6 [28] using manual thresholds when automatic thresholding failed, ambient RNA was removed using DecontX 1.12 [29] and cells estimated to contain >50% ambient RNA were removed, and doublets were identified using DoubletFinder 2.0.3 [30] and removed. The remaining cells were then filtered to retain only those with > 200 genes, 500-80,000 UMIs, < 10-20% of UMIs from genes encoded by the mitochondrial genome (sample dependent based on UMI distributions), < 5% of UMIs derived from HBB, < 20,000 UMIs from antibody-derived tags (ADTs), and >100-2,750 UMIs from antibody derived tags (sample dependent based on UMI distribution). Filtered cells were modeled in latent space using TotalVI 0.18.0 [31] to create a joint embedding derived from both RNA and ADT expression data, corrected for batch effects, mitochondrial proportion, and cell cycle. Scanpy 1.8.2 [32] was used to cluster the data in latent space using the Leiden algorithm [33], and marker genes were identified in latent space using TotalVI. Clusters were annotated using clustifyr 1.9.1 [34] and the leukemic/normal bone marrow reference dataset presented in [35]. Scanpy and Seurat 4.1.1 [36] were then used to generate UMAP projections from the TotalVI embeddings and perform exploratory analysis, data visualization, etc.

### Variant calling on AML panel pulldowns

Single-cell alignments from Star-solo as well as valid barcodes were input into cellsnp-lite [20] in bulk calling mode to generate a starting list of variants. Variants were filtered on 1 of 2 criteria: A) if the variant was in a gene not in the pulldown list we selected variants that met 3 criteria; 1) an alternate allele depth of 50 2) a reference allele depth of 50 3) an alternate allele ratio of less than 80%, B) if the variant was in the pulldown list required both an alternate depth of 10 and an alternate allele ratio of less than 90%. This filtered variant list was then utilized to call variants on single cells with Cellsnp-lite in single-cell calling mode. The resulting variant matrices were used as input into Seurat to visualize mutational profiles across cells.

### Variant calling on mitochondrial panel pulldowns

Mitochondrial variants were called using Cellsnp-lite [20]. The resulting matrices were then filtered with MQUAD [19] to identify informative variants for clonal lineage reconstruction. These filtered variants were then used to assign clones with vireoSNP[24] as described in [19] and visualized with Seurat.

## Acknowledgments

We thank Clay Smith, Kent Riemondy, Shashan Pei, William Showers, Sarah Staggs, Abbigayl Burtis, Stephanie Gipson, and Maura Gasparetto. for their invaluable feedback during the development of these techniques and preparation of the resulting manuscript.

